# Temperature modulates resting posture, microhabitat use, and day nesting in chimpanzees in a cool montane forest

**DOI:** 10.64898/2026.08.18.745634

**Authors:** Hassan Al-Razi, Fidele Muhayeyezu, Felix Mulindahabi, Protais Niyigaba, Drew Arthur Bantlin, Beth A. Kaplin, Shane K. Maloney, Cyril C. Grueter

## Abstract

Behavioral thermoregulation allows animals to reduce the physiological costs associated with variation in the thermal environment, yet little is known about how chimpanzees respond behaviorally to the cool and humid conditions of tropical montane forests. We examined how microclimatic conditions influenced resting posture, sun–shade use, substrate use, vertical position within the canopy, and day-nest use in eastern chimpanzees (*Pan troglodytes schweinfurthii*) in Nyungwe National Park, Rwanda. From March 2024 to February 2025, we collected 5,661 individual behavioral records during 486 contact hours and paired behavioral observations with measurements of ambient temperature, water vapor pressure, wind speed, and rainfall. Bayesian mixed models showed that temperature was the most consistent climatic correlate of resting behavior. Under cooler conditions, chimpanzees were more likely to adopt heat-conserving postures, particularly curled and huddled postures, and to use day nests. As temperature increased, chimpanzees increased their use of shade, shifted from arboreal to terrestrial resting, and used lower canopy strata more frequently. Higher wind speed was also associated with reduced use of the upper canopy, whereas rainfall, water vapor pressure, and fruit availability generally showed weaker or behavior-specific associations. These findings indicate that chimpanzees in a cool montane forest flexibly adjust posture, solar exposure, substrate use, vertical position, and day-nest use in relation to local thermal conditions. Resting behaviour may therefore provide an important context for behavioral thermoregulation, extending our understanding of chimpanzee thermal ecology beyond the warmer lowland and savanna habitats that have received most previous attention.

## 1 Introduction

Rest encompasses periods of behavioral inactivity that serve multiple adaptive functions and is an important component of the activity pattern of an animal. Rather than being a reflection simply of time that is not allocated to an active behavior, rest time is essential for physiological recuperation, digestion, predator avoidance, and thermoregulation (Herbers, 1981). As endotherms, mammals, including primates, must maintain a stable body temperature in the face of environmental variability. Rest provides a key behavioral context in which individuals can alter their heat exchange with the environment because the production of metabolic heat is minimal when an animal is at rest, and the opportunity to adjust posture or position can alter the exposure to external conditions (Terrien et al., 2011; Korstjens et al., 2010). Consequently, variation in rest behavior can provide insight into how an animal balances energetic demands with thermal constraints, as time budgets reflect fundamental trade-offs among foraging, survival, and environmental conditions (Korstjens et al., 2010; Shukla et al., 2021; Kluiver et al., 2022). Climatic factors play a central role in the shaping of these patterns, as animals modify rest behaviour and habitat use to mitigate thermal stress, either via the conservation of heat under cold conditions or the minimisation of heat gain in hot environments (Stelzner, 1988; Roberts & Dunbar, 1991; Hill, 1999).

During periods of inactivity, animals, including primates, employ a range of behavioral strategies that contribute to thermoregulation. These strategies, such as the adoption of specific postures, the selection of a favorable microhabitat, or the use of particular vertical strata, can reduce the metabolic requirements or enhance heat dissipation, and thereby reduce the physiological cost of thermoregulation under varying environmental conditions (Thompson & Hermann, 2024). Among these, the adoption of a certain posture is a key and well-documented thermoregulatory behaviour in primates (Paterson, 1981; Stelzner & Hausfater, 1986; Bicca-Marques & Calegaro-Marques, 1998; Terrien et al., 2011). In particular, as ambient temperature and humidity increase, non-human primates tend to shift towards a posture that facilitates heat dissipation (Bicca-Marques & Calegaro-Marques, 1998; Gestich et al., 2014; Lopes & Bicca-Marques, 2017; Brividoro et al., 2023; Chen-Kraus et al., 2023), whereas under cooler conditions they adopt a posture that is conducive to heat-conservation (Grueter et al., 2013). Social behaviors can also contribute to posture-based thermoregulation; for example, huddling reduces heat loss and helps individuals maintain body temperature under cold conditions (Bicca-Marques & Calegaro-Marques, 1998; Li et al., 2010; Gestich et al., 2014), and allogrooming can help to maintain the insulative capacity of the pelt (McFarland et al., 2016). Individual vervet monkeys that had more social connections, and so presumably received more allogrooming, maintained a higher body temperature under cold conditions than did those that had fewer social connections (McFarland et al., 2015; Henzi et al., 2017).

Microhabitat selection is another important behavioral strategy through which animals can avoid thermal stress. The selection of shaded areas under hot conditions and sun-exposed locations in colder conditions can modulate heat gain and loss (Carrascal et al., 2001; Brown & Downs, 2002; Smith, 1983; Kelley et al., 2016; Campbell et al., 2018a; McFarland et al., 2020). In primates, microhabitat choices are often closely linked to the resting location, as individuals choose terrestrial or arboreal sites that differ in thermal properties and microclimatic conditions (Eppley et al., 2022; Souza-Alves et al., 2019; Mekonnen et al., 2018; Fruth & McGrew, 1998). Ground-level environments can provide shaded, thermally buffered conditions, whereas arboreal positions can increase exposure to solar radiation and /or convective cooling depending on canopy structure and wind, allowing a primate to exploit vertical heterogeneity in the thermal environment (Samson & Hunt, 2012; Thompson et al., 2016; Palestino-Sánchez et al., 2025a; Mekonnen et al., 2018; Mourthé et al., 2007). This vertical heterogeneity also extends within an arboreal setting, where different canopy strata vary in microclimatic conditions such as temperature, solar exposure, and wind. Lower strata are typically more sheltered and thermally stable, whereas higher canopy layers are more exposed to solar radiation and airflow, allowing individuals to further modulate heat gain and heat loss according to the prevailing environmental conditions (Palestino-Sánchez et al., 2025a, 2025b; Li, 2007; Mekonnen et al., 2018). Together, the selection of a resting location and canopy stratum can enable a primate to fine-tune heat exchange with the environment across a vertical gradient.

Variation in rest posture and substratum use has been documented across chimpanzee populations, and reflects behavioral flexibility under local environmental conditions. At Bossou, chimpanzees were more terrestrial under hotter and drier conditions than under cooler wet conditions, suggesting that ground use may provide access to favorable microclimates (Takemoto, 2004). In Budongo, temperature was positively associated with rest and negatively associated with feeding. Chimpanzees also increased terrestriality and reduced their use of sun-exposed areas during periods of high temperature (Kosheleff & Anderson, 2009). Similar thermoregulatory flexibility has been reported in captive chimpanzees through shade use (Duncan & Pillay, 2013), and in savanna chimpanzees at Fongoli through cave use during hot conditions (Pruetz, 2007). Together, these studies show that chimpanzees adjust their use of substratum, exposure, and rest behavior in response to local thermal environments.

In this context, day nesting may be another important, but understudied, form of behavioral thermoregulation in chimpanzees. Like other great apes, chimpanzees build nests not only for night-time sleep but also for daytime rest. However, day nests are usually used for a short period and have received less research attention than have night nests (Fruth & Hohmann, 1996; Anderson, 2000; Stewart, 2011; Koops et al., 2012; Stewart et al., 2018; McGrew, 2021). During rest, day nests may offer several benefits, including support for the body, comfort, protection from uncomfortable substrate, and buffering from local weather conditions (Boesch, 1995; Fruth & Hohmann, 1996; Brownlow et al., 2001). By the accumulation of vegetation beneath the body, a nest may also create a more insulated resting surface, consistent with evidence that chimpanzee nests can provide both structural support and thermal insulation, and that nest structure can vary with local weather conditions (Fruth & Hohmann, 1996; Stewart et al., 2018; Al-Razi et al., 2026). Under cool conditions, day nests may therefore help chimpanzees to conserve heat while they rest, especially during inactive periods when they produce less metabolic heat. The value of an arboreal day nest, however, may also depend on wind, which increases heat loss through convection. Nests that are built above the ground or in exposed parts of the canopy may leave resting chimpanzees more vulnerable to cooling in windy conditions. In line with this idea, Boesch (1995) reported that chimpanzees in Taï National Park built day nests more frequently on the ground during the Harmattan, a dry and windy period, than at other times. This suggests that, when conditions are windy, chimpanzees may avoid arboreal resting sites that are exposed and instead use ground nests to reduce heat loss from wind exposure.

However, most studies of rest behavior in chimpanzees have been conducted in relatively warm lowland forests, in captivity, or in savanna-mosaic habitats, where the daytime thermal challenge is usually a heat load rather than cool conditions. In contrast, tropical montane forests are characterized by lower temperatures and greater precipitation, wind, and cloud cover compared to lowland forest (Chapman et al., 2016; Bruijnzeel et al., 2010). Little is known about how chimpanzees adjust their resting behavior during prolonged exposure to cool, wet and thermally variable conditions, particularly at high elevations. To address this gap, we investigated resting posture and substratum use in Nyungwe National Park, one of the highest-elevation habitats occupied by chimpanzees, ranging from approximately 1,600 to 2,950 m a.s.l. Nyungwe is characterized by relatively low temperatures, high rainfall and frequent wind exposure (Gross-Camp & Kaplin, 2005; Williamson et al., 2013; Nyirambangutse et al., 2017; Green et al., 2020a). Its thermal environment therefore contrasts with the warmer and more open habitats in which most previous studies of chimpanzee resting behavior have been conducted. The combined effects of low ambient temperatures, rainfall and wind may increase heat loss and expose chimpanzees to prolonged thermal challenges.

In this study, we examined how temperature, humidity, rainfall, and wind speed influence the posture and microhabitat used during rest, and the use of a day-nest, in eastern chimpanzees (*Pan troglodytes schweinfurthii*) in Nyungwe National Park. Specifically, we tested whether chimpanzees adjust these behaviors as thermoregulatory strategies under cool montane forest conditions. By integrating fine-scale microclimatic measurements with detailed behavioral observations, we assessed how individuals respond to short-term thermal variation. We focused particularly on rest behavior because rest posture and rest-site or microhabitat choice are widely recognized as important components of behavioral thermoregulation in mammals, including primates (Stelzner & Hausfater, 1986; Terrien et al., 2011; Gestich et al., 2014; Lopes & Bicca-Marques, 2017). In contrast, feeding and locomotion are often strongly constrained by food availability, food distribution, travel costs, and social factors, making thermoregulatory effects more difficult to isolate in those behavioral contexts (Isbell & Young, 1993; Chapman et al., 1995). Rest, therefore, provides a more appropriate behavioral context to assess whether posture, microhabitat use, and use of a day-nest are modified by the thermal environment. Accordingly, we tested the following hypotheses related to rest behavior:

H1. At lower temperatures, chimpanzees will increase their use of heat-conserving postures, such as curled or huddled postures. The opposite pattern is expected at higher temperatures. Higher wind speed is also expected to increase the use of heat-conserving postures because of increased convective heat loss.

H2. At higher temperatures, chimpanzees will increase their use of shaded microhabitats and reduce the use of sun-exposed microhabitats. Higher humidity is also expected to promote shade use because reduced evaporative cooling efficiency may increase thermal discomfort. Conversely, lower temperature is expected to lead to rest in more sun-exposed positions.

H3. Chimpanzees will adjust their use of the substratum and canopy stratum in response to climatic conditions. At lower temperatures, individuals are expected to use higher canopy strata more frequently because these positions provide greater solar exposure. Under windier conditions, individuals are expected to use the ground and lower canopy strata more frequently, where vegetation structure may provide shelter from the wind.

H4. Rainfall will influence the selection of microhabitat for rest. During rainfall, chimpanzees are expected to reduce ground rest and increase arboreal rest. Within trees, individuals are expected to preferentially use lower strata, where canopy cover and vegetation structure may reduce direct exposure to rain.

H5. Chimpanzees are expected to use day nests more frequently under cooler conditions. In contrast, the use of a day nest is expected to decrease under windy conditions, when arboreal nests may increase exposure to convective heat loss.

## 2 Methods

### 2.1 Ethical Note

This non-invasive observational study was approved by the Animal Ethics Committee of The University of Western Australia (approval no. F18979-05, 26 October 2023). Research permission in Rwanda was granted by the National Council for Science and Technology (permit no. NCST/482/0086/2024) and the Rwanda Development Board, with access and research consent provided by Nyungwe Management Company/African Parks. Observations involved no handling, trapping, experimentation, habitat alteration, or collection of biological samples. Researchers maintained a minimum distance of 10 m from the chimpanzees to minimize disturbance and pathogen-transmission risk. All fieldwork complied with Nyungwe National Park regulations, applicable Rwandan research requirements, and the American Society of Primatologists’ Principles for the Ethical Treatment of Non-Human Primates.

### 2.2 Study Area and Species

Nyungwe National Park (NNP), located in southwestern Rwanda (2°17′–2°50′S, 29°07′– 29°26′E), encompasses approximately 1,013 km² of tropical montane forest within the Albertine Rift. The park spans a pronounced elevational gradient (c. 1,600–2,950 m a.s.l.) and has a cool, humid environment that is characterized by relatively stable temperatures and high precipitation throughout the year. Rainfall occurs in two main wet seasons (September– November and January–March) that are interspersed with comparatively drier periods from April to August and in December, although the timing and intensity of rainfall have considerable interannual variability (Sun et al., 1996; Nyirambangutse et al., 2017; Matthews et al., 2019). NNP forms part of the contiguous Nyungwe–Kibira forest block, one of the largest remaining montane forest systems in Central Africa (Plumptre et al., 2002). This ecosystem supports high biodiversity, including more than 80 mammal species, numerous Albertine Rift endemics, and at least 12 species of non-human primate. The vegetation is structurally and compositionally heterogeneous, comprising a mosaic of primary and secondary forest, bamboo zones, swamp habitats, and montane grasslands. Local habitat variation is strongly influenced by elevation, canopy structure, and moisture gradients (Plumptre et al., 2010).

The study focused on the well-habituated Mayebe chimpanzee (*Pan troglodytes schweinfurthii*) community. During the study period, the community consisted of approximately 67 individuals, including adult and subadult males and females, juveniles, and infants. The community occupies a home range between Uwinka and Mount Bigugu that covers approximately 38–48 km² depending on the estimation method (Green et al., 2020a, 2020b). Notably, this range includes the upper elevational limits of chimpanzee distribution, reaching nearly 2,950 m above sea level. As in other chimpanzee populations, the Mayebe community exhibits a multimale–multifemale social organization and a fission–fusion dynamic, with frequent changes in subgroup size and composition (Matthews et al., 2021). These social and ecological characteristics, combined with the unique high-elevation environment, make this population particularly suitable to investigate behavioral responses to montane climatic conditions (Al-Razi et al., 2026).

### 2.3 Behavioural Data Collection

We conducted behavioral observations between March 2024 and February 2025 with support from two trained field assistants, sampling on approximately eight days each month. Over the study period, we accumulated 486 contact hours, which comprised 1,933 party-level scans and 5,661 individual records using instantaneous scan sampling methods (Altmann, 1974). The field conditions in Nyungwe, particularly the rugged terrain, dense forest structure, and large ranging pattern of chimpanzees, limited our ability to carry out continuous focal follows. To address this, we implemented a structured scan-based approach at the party level (Matthews et al., 2021). Within each sampling bout, we systematically recorded visible individuals in a consistent directional order, which allowed us to standardize observations, prevent duplicate sampling within scans, and reduce potential observer bias associated with conspicuous individuals or behaviors (Fashing, 2001; Mekonnen et al., 2018; Bateson & Martin, 2021). We initiated daily follows by relocating the nesting site that was used the previous night, which enabled us to monitor chimpanzee parties from early morning departure through to evening nest construction. This approach facilitated continuous, full-day behavioral coverage whenever possible. In cases where the nest site could not be identified, we located chimpanzees by tracking vocal cues, recent feeding traces, and fresh signs of movement. We maintained contact with the same party for as long as possible; when a party was lost, we attempted re-establishment, and if unsuccessful, we continued sampling on a newly encountered group (Gilby et al., 2010).

To examine the influence of climatic variables on chimpanzee activity, we classified behavior into four primary categories: feeding, rest, locomotion, and grooming (van Lawick-Goodall, 1968; Kosheleff & Anderson, 2009). Other social behaviors, such as play and aggression, were excluded due to their low frequency and the limited ability to robustly assess their relationships with environmental variables. Feeding included all food processing and ingestion behaviors. Rest was defined as any inactive state, regardless of posture (e.g., sitting, lying, or stationary standing). Locomotion encompassed all forms of movement, including walking and climbing, while grooming included both self-directed and social grooming.

Although we recorded all major activity types, we restricted the detailed analyses of posture and microhabitat use to those individuals whose primary activity was rest. Rest provides the clearest context to assess body posture and exposure to local environmental conditions, because individuals are inactive and less affected by the immediate demands of feeding, moving, or social interaction (Gestich et al., 2014). During scan sampling, we therefore recorded body posture for all individuals that were considered at rest, and classified their posture following established schemes (Paterson, 1981; Grueter et al., 2013; Gestich et al., 2014). Posture was categorized as: (i) curled, with limbs flexed close to the body and the back bent forward; (ii) sitting, where the individual rests on its haunches with feet positioned either toward or away from the body midline; (iii) lying, where the body and head are supported against a substrate with limbs flexed; (iv) huddling, in which two or more individuals rested in close body contact; and (v) spread, characterized by limbs extended away from the body, usually in a horizontal or splayed position, resulting in a more open body posture. The spread posture was excluded from subsequent analyses due to insufficient observations. To characterize microhabitat conditions during rest, we recorded whether individuals were positioned on the ground or in a tree, and whether they were exposed to direct sunlight or in a shaded condition. For arboreal rest, we further classified the vertical position within the canopy as low, mid, or top strata following established approaches (Palestino-Sánchez et al., 2025b). In addition, we recorded whether each resting individual used a day nest. Individuals were classified into five age–sex groups: adult male, adult female, subadult male, subadult female, and juvenile, following standard criteria for eastern chimpanzees (van Lawick-Goodall, 1968). Infants were omitted from all analyses.

### 2.4 Weather Data Collection

To quantify the influence of climatic variables on posture and microhabitat use, we recorded fine-scale microclimatic conditions during each behavioral scan, including ambient temperature, relative humidity, wind speed, and rainfall. Rainfall was treated as a binary variable (present / absent). During each scan, we measured ambient temperature and wind speed using a handheld anemometer (Extech 45158 Rotary Vane), while relative humidity was recorded with a handheld heat-stress meter (Extech HT30 WBGT Meter). To ensure that our measurements reflected the condition that was experienced by the chimpanzees, we matched the sampling location to the individuals’ immediate exposure. To approximate the wind condition that was experienced by an arboreal individual, we accounted for vertical position by having a field assistant move to a nearby elevated point or slope that approximated the height of the focal individual in the canopy. All measurements were taken within 25 m of the party location. Wind speed was measured with the anemometer oriented into the prevailing wind and positioned away from surrounding vegetation or other obstructions to minimize turbulence and measurement bias.

Relative humidity is strongly temperature-dependent, and so, for analysis, we converted the relative humidity to water vapor pressure (WVP), which provides a measure of absolute humidity and helps to reduce collinearity with air temperature and improve the stability of statistical analyses. Following Barenbrug (1974), we calculated the water vapor pressure as:

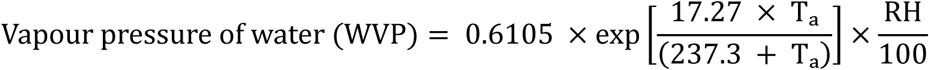

Where:

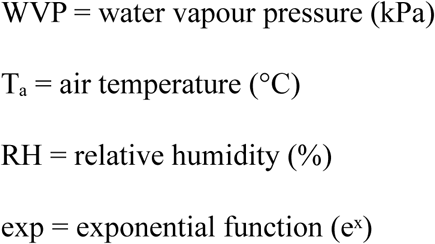

### 2.5 Fruit Availability Index (FAI)

The availability of fruit can modulate positional and postural behavior in primates (Dasilva, 1993; Matsuda et al., 2017; Zheng et al., 2021). Therefore, to assess the potential relationship between fruit availability, rest posture, and microhabitat use, we monitored 19 tree species that were identified as important chimpanzee food resources during the study period. Within the home range of the chimpanzees, we sampled a total of 620 individual trees that belonged to these species across 17 selected trails, with each species represented by more than 15 individuals. Only trees that had a diameter at breast height (DBH) ≥ 10 cm and a crown that was clearly visible from the trail were monitored to ensure a reliable assessment of fruiting. For each tree, we scored the proportion of fruit that was present each month using a five-point ordinal scale: 0 (<1%), 1 (1–25%), 2 (26–50%), 3 (51–75%), and 4 (76–100%). We calculated a monthly fruit availability index (FAI) by weighting the fruiting score by tree size (DBH) and summed values across all monitored trees for each month, using the following equation:

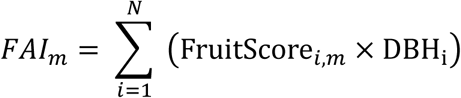

Where FAI represents fruit availability in month *m*, FruitScore is the fruit score of tree *i* in month *m*, DBH is the diameter at breast height of tree *i*, and *N* is the total number of monitored trees. Further details on the establishment of phenology routes and comparable data-collection protocols are described in Sun et al. (1996) and Dugger et al. (2025).

### 2.6 Statistical analysis

All analyses were conducted in R version 4.3.0 (R Core Team, 2023). Data handling and preprocessing were performed using the *tidyverse* package (Wickham et al., 2019), which was used for data cleaning, transformation, aggregation, and reshaping. Behavioral and environmental datasets were cleaned and filtered prior to analysis to ensure consistency across models. Because environmental variables were measured at the scan level, behavioral responses were also aggregated to the scan level before modelling. For each scan, we calculated the number of resting chimpanzees exhibiting each behavioral state or microhabitat-use category. This ensured that the analytical unit corresponded to the scale at which environmental predictors were measured and reduced pseudo-replication arising from multiple individuals being observed under the same environmental conditions. To improve model convergence, facilitate comparison of effect sizes among predictors, and reduce potential scaling issues, continuous predictors were standardized to a mean of 0 and standard deviation of 1. Ambient temperature, water vapor pressure (WVP), wind speed, and fruit availability index (FAI) were standardized within the analysis dataset. Because climatic predictors may be correlated, we assessed multicollinearity using variance inflation factors (VIFs) with the *car* package (Fox & Weisberg, 2019), with results additionally checked using the *performance* package (Lüdecke et al., 2021). All environmental predictors had VIF values < 3, indicating low multicollinearity and supporting their inclusion within the same models.

We fitted Bayesian generalized linear mixed models using the brms package (Bürkner, 2017), which provides an interface to Stan for Bayesian multilevel modelling. Weakly informative priors were used throughout. Population-level regression coefficients were assigned Normal(0, 1) priors on the log-odds scale. Additional weakly informative priors were specified for intercepts and group-level standard deviations where appropriate; otherwise, default brms priors were retained (Table S1). Models were fitted using four Markov chain Monte Carlo (MCMC) chains with 4,000 iterations per chain, including 2,000 warm-up iterations, yielding 8,000 post-warm-up posterior draws. Day identity was included as a random intercept in all models to account for repeated sampling within days and unmeasured day-level heterogeneity. Because behavioral observations were obtained at 15-min intervals, the behavioral composition of successive scans could show short-term temporal dependence. We therefore incorporated the behavioral composition of the immediately preceding scan within the same sampling day as a temporal-control predictor. For binary responses, this variable was the proportion of individuals exhibiting the focal behavior in the preceding scan. For multinomial responses, the proportions of each non-reference category in the preceding scan were included as temporal-control predictors. The reference-category proportion was omitted because it could be calculated from the remaining proportions. Scans were ordered chronologically within each day, and the first scan of each day was excluded from the corresponding model because no previous-scan composition was available. These lagged terms were included to account for first-order temporal persistence in scan-level behavioral composition rather than to represent individual behavioral continuity.

For the resting-posture model, we aggregated observations within each scan and modelled the numbers of chimpanzees sitting, lying, curled, or huddled using a multinomial GLMM with a logit link, with sitting as the reference category. To account for short-term temporal dependence, we also included the proportions of chimpanzees that were lying, curled, or huddled in the previous scan.

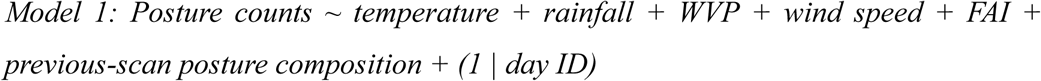

To assess the effect of microclimatic conditions on sun-shade use, analyses were restricted to scans conducted under sunny conditions, when the sun was not obscured by cloud. For each scan, the number of chimpanzees resting in shade was modelled relative to the total number of shade and sun observations using a binomial model with a logit link. The proportion of individuals using shade in the immediately preceding scan was included as the temporal-control variable.

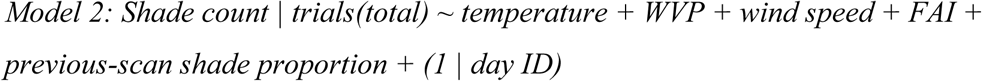

To examine how microclimatic conditions influenced substratum use during resting, the number of chimpanzees resting in trees within each scan was modelled relative to the total number resting either in trees or on the ground using a binomial model with a logit link. The proportion of chimpanzees using trees in the previous scan was included to account for short-term temporal dependence in tree use.

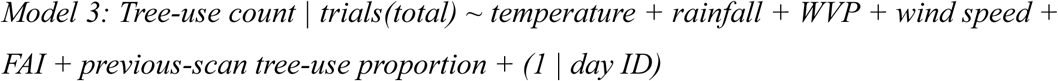

To examine vertical space use within trees, analyses were restricted to arboreal resting observations. The numbers of chimpanzees occupying the low, middle, and upper canopy strata within each scan were modelled jointly using a multinomial GLMM with a logit link, with the low canopy as the reference category. The proportions of chimpanzees using the middle and upper canopy in the previous scan were included to account for short-term temporal dependence.

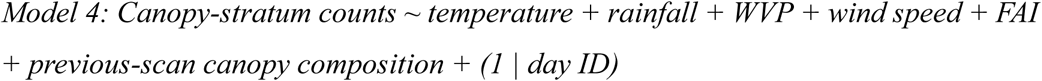

To examine how microclimatic conditions influenced day-nest use, the number of chimpanzees resting in day nests within each scan was modelled relative to the total number of nest and non-nest resting observations using a binomial model with a logit link. The proportion of chimpanzees using a day nest in the previous scan was included to account for short-term temporal dependence.

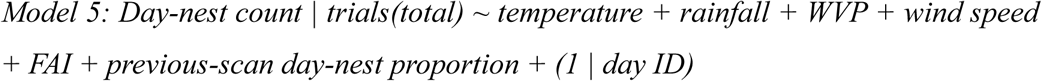

Because rainfall could modify the effect of temperature in cool montane forests, we initially fitted models containing a temperature × rainfall interaction. However, there was little posterior support for these interaction effects, and their 95% credible intervals overlapped zero. The interaction terms were therefore removed from the final models to improve model parsimony and interpretability.

Model convergence was assessed using R^, effective sample sizes (ESS), and divergent transitions. R^ values close to 1.00, adequate ESS, and no divergent transitions were considered evidence of satisfactory convergence. Posterior estimates are reported with 95% credible intervals (CrIs), with effects considered strongly supported when CrIs did not include zero. Predicted probabilities were obtained using posterior_epred, with other predictors held at their means or reference levels and day-level random effects excluded to show population-level relationships. Temperature was back-transformed to °C for plotting. Original individual observations were displayed as jittered 0/1 points, whereas fitted relationships and 95% CrIs were derived from the scan-level models.

## 3 Results

The resting posture of chimpanzees was strongly associated with temperature (Table 1; Fig. 1A). Relative to sitting, the probabilities of lying, curling, and huddling all decreased with increasing temperature (Fig. 1B), with the strongest effect observed for huddling (β = −2.05, 95% CrI [−2.59, −1.54]), followed by curled posture (β = −0.82, [−1.04, −0.62]) and lying (β = −0.37, [−0.62, −0.13]). Water vapor pressure was also negatively associated with huddling (β = −0.54, [−1.04, −0.06]), whereas rainfall, wind speed, and fruit availability showed no clear associations with posture. Previous-scan posture composition indicated short-term temporal dependence. Lying was more likely when a greater proportion of chimpanzees had been lying in the previous scan (β = 2.10, 95% CrI [1.55, 2.63]) and less likely when more had been huddling (β = −1.06, [−2.13, −0.06]). Curled posture was more likely following scans with higher proportions of curled (β = 0.92, [0.49, 1.36]) or huddling individuals (β = 1.21, [0.57, 1.85]). No previous-scan posture measure showed a clear association with current huddling. Day-level variation was greatest for huddling (SD = 1.94, 95% CrI [1.27, 2.85]), followed by lying (SD = 1.31, [0.93, 1.79]) and curled posture (SD = 0.42, [0.05, 0.77]).

**Figure 1.**
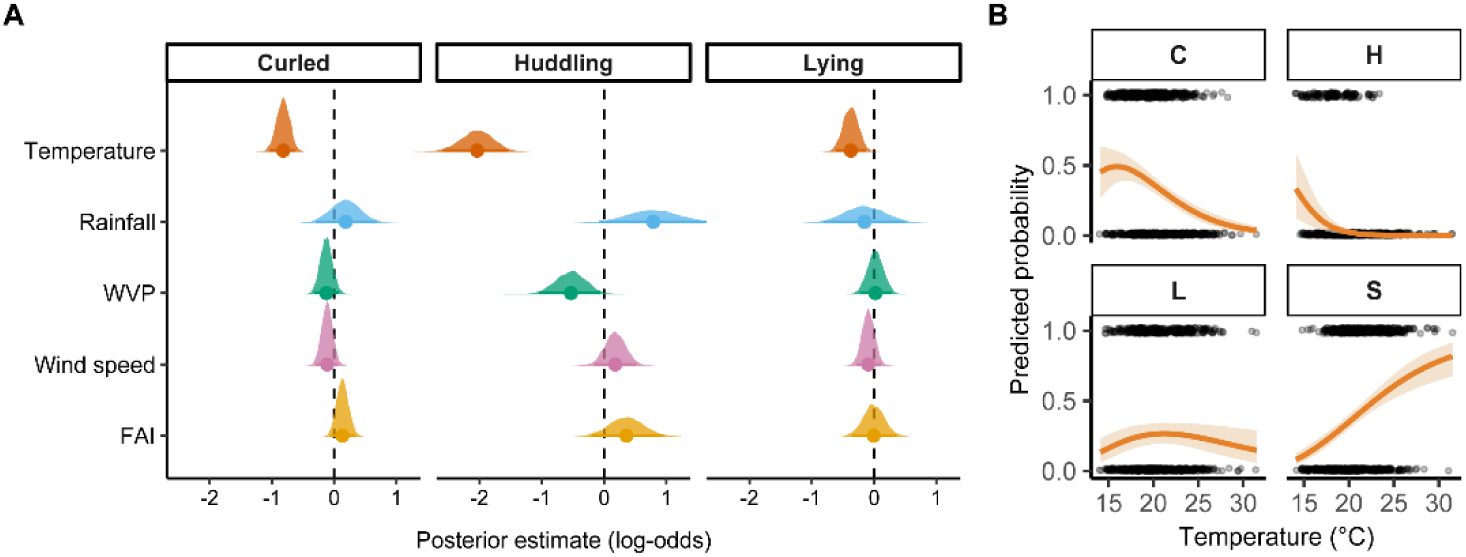
Effects of microclimatic variables on resting posture in chimpanzees. (A) Posterior estimates from the Bayesian multinomial GLMM for the effects of temperature, rainfall, water vapour pressure (WVP), wind speed, and fruit availability index (FAI) on curled, huddled, and lying postures relative to sitting. Points indicate posterior means, shaded distributions show posterior uncertainty, and the dashed vertical line indicates no effect. Negative estimates indicate lower odds relative to sitting, and positive estimates indicate higher odds. (B) Predicted probabilities of each resting posture across the observed temperature range. Lines show posterior mean predictions and shaded bands represent 95% credible intervals; points show observed individual posture records. Posture categories are C = curled, H = huddled, L = lying, and S = sitting.

**Table 1.** Posterior estimates from a Bayesian multinomial generalized linear mixed model (GLMM) examining the effects of microclimatic variables, fruit availability, and previous resting posture on chimpanzee resting posture. Estimates are presented with posterior standard errors and 95% credible intervals (CrI). Day was included as a random intercept (81 levels), with random effects reported as standard deviations (SD). Model convergence was satisfactory, with all R^ values equal to 1.00 and adequate effective sample sizes.

| Response category | Predictor | Estimate | Est. error | Lower 95% CrI | Upper 95% CrI | $\hat{R}$ |
| --- | --- | --- | --- | --- | --- | --- |
| Lying | Intercept | -0.88 | 0.24 | **-1.35** | **-0.40** | 1.00 |
|  | Temperature | -0.37 | 0.12 | **-0.62** | **-0.13** | 1.00 |
|  | Rainfall (yes) | -0.15 | 0.34 | -0.81 | 0.51 | 1.00 |
|  | Water vapour pressure | 0.02 | 0.13 | -0.23 | 0.28 | 1.00 |
|  | Wind speed | -0.09 | 0.10 | -0.29 | 0.11 | 1.00 |
|  | Fruit availability index | -0.00 | 0.18 | -0.35 | 0.36 | 1.00 |
|  | Previous posture: lying | 2.10 | 0.27 | **1.55** | **2.63** | 1.00 |
|  | Previous posture: curled | 0.12 | 0.27 | -0.40 | 0.64 | 1.00 |
|  | Previous posture: huddling | -1.06 | 0.53 | **-2.13** | **-0.06** | 1.00 |
| Curled | Intercept | -0.51 | 0.16 | **-0.83** | **-0.19** | 1.00 |
|  | Temperature | -0.82 | 0.11 | **-1.04** | **-0.62** | 1.00 |
|  | Rainfall (yes) | 0.18 | 0.25 | -0.30 | 0.68 | 1.00 |
|  | Water vapour pressure | -0.12 | 0.10 | -0.32 | 0.08 | 1.00 |
|  | Wind speed | -0.11 | 0.09 | -0.29 | 0.06 | 1.00 |
|  | Fruit availability index | 0.14 | 0.10 | -0.05 | 0.33 | 1.00 |
|  | Previous posture: lying | -0.15 | 0.30 | -0.76 | 0.43 | 1.00 |
|  | Previous posture: curled | 0.92 | 0.22 | **0.49** | **1.36** | 1.00 |
|  | Previous posture: huddling | 1.21 | 0.33 | **0.57** | **1.85** | 1.00 |
| Huddling | Intercept | -3.14 | 0.49 | **-4.20** | **-2.29** | 1.00 |
|  | Temperature | -2.05 | 0.26 | **-2.59** | **-1.54** | 1.00 |
|  | Rainfall (yes) | 0.79 | 0.44 | -0.08 | 1.68 | 1.00 |
|  | Water vapour pressure | -0.54 | 0.25 | **-1.04** | **-0.06** | 1.00 |
|  | Wind speed | 0.18 | 0.17 | -0.16 | 0.51 | 1.00 |
|  | Fruit availability index | 0.36 | 0.29 | -0.20 | 0.95 | 1.00 |
|  | Previous posture: lying | -0.11 | 0.49 | -1.09 | 0.84 | 1.00 |
|  | Previous posture: curled | -0.25 | 0.40 | -1.03 | 0.53 | 1.00 |
|  | Previous posture: huddling | 0.43 | 0.48 | -0.53 | 1.35 | 1.00 |
| Random effects | SD (day): lying | 1.31 | 0.22 | **0.93** | **1.79** | 1.00 |
|  | SD (day): curled | 0.42 | 0.18 | **0.05** | **0.77** | 1.00 |
|  | SD (day): huddling | 1.94 | 0.40 | **1.27** | **2.85** | 1.00 |

Shade use increased strongly with temperature (β = 1.85, 95% CrI [1.31, 2.44]; Table 2, Fig. 2A). Chimpanzees were therefore more likely to rest in shade under warmer conditions (Fig. 2B). Water vapor pressure, wind speed, and fruit availability showed no clear associations with shade use. Previous-scan shade use was also strongly associated with current shade use (β = 2.01, 95% CrI [1.21, 2.82]), indicating short-term temporal persistence in shade use. Day-level variation was substantial (SD = 2.77, 95% CrI [1.76, 4.19]).

**Figure 2.**
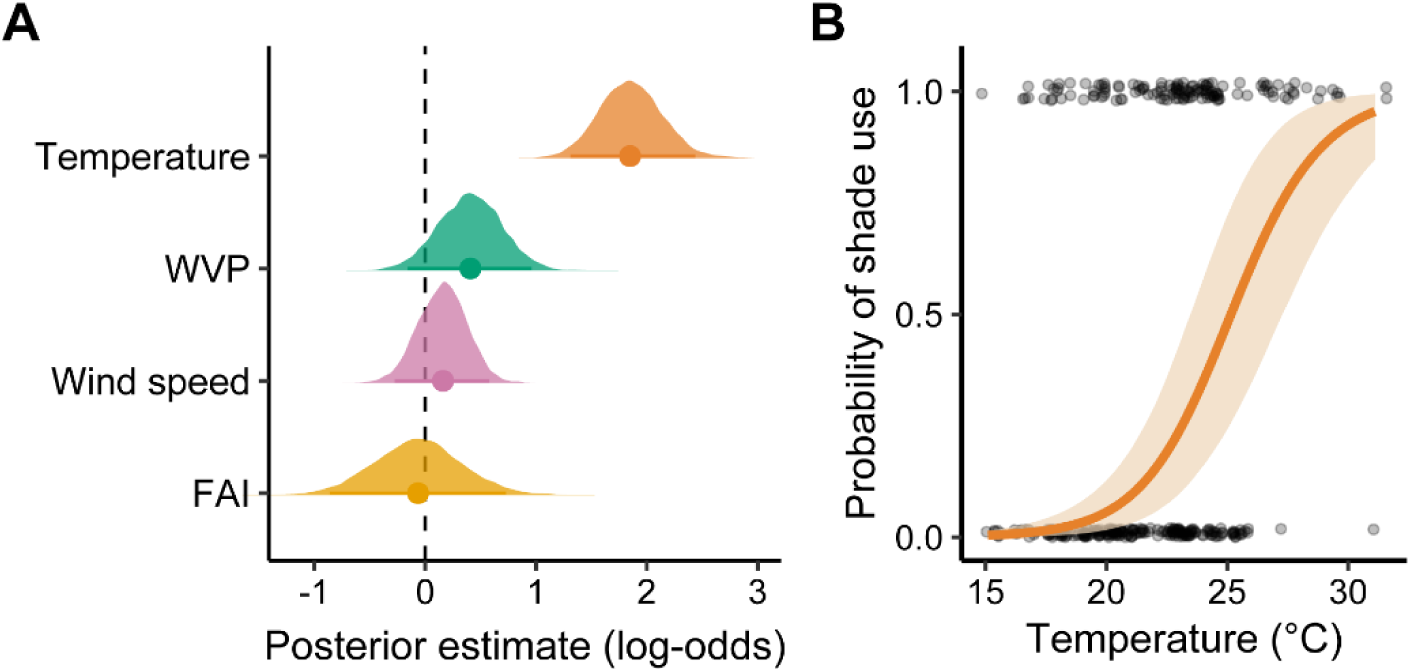
Effects of microclimatic variables on shade use during resting. (A) Posterior estimates from the Bayesian binomial GLMM for the effects of temperature, water vapour pressure (WVP), wind speed, and fruit availability index (FAI) on shade use. Points indicate posterior means, shaded distributions show posterior uncertainty, and the dashed vertical line indicates no effect. Positive estimates indicate increased odds of shade use, whereas negative estimates indicate decreased odds. (B) Predicted probability of shade use across the observed temperature range. The line shows the posterior mean prediction and the shaded band represents the 95% credible interval. Points show observed individual records, with shade coded as 1 and sun exposure as 0.

**Table 2.**
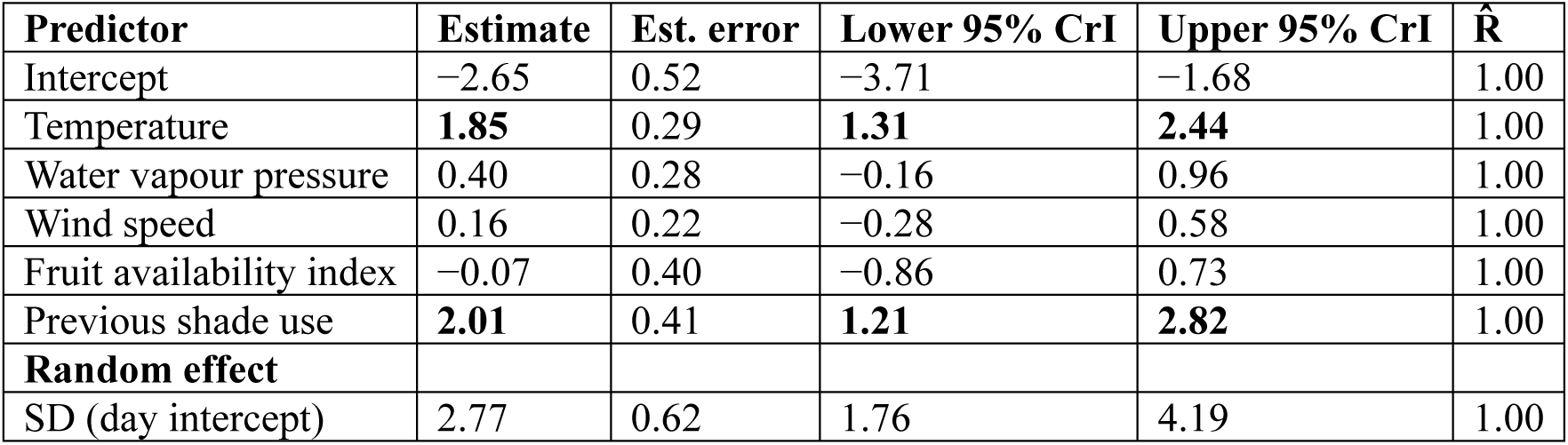
Posterior estimates from a Bayesian binomial generalized linear mixed model (GLMM) examining shade use by chimpanzees during resting in relation to microclimatic variables, fruit availability, and temporal dependence. Estimates are presented on the log-odds scale, with positive values indicating an increased probability of shade use. The lagged term (previous shade use) accounts for temporal dependence between consecutive observations. Day was included as a random intercept, with the random effect reported as the standard deviation (SD). Model convergence was satisfactory based on R^ values and effective sample sizes.

Tree use declined with increasing temperature (β = −0.70, 95% CrI [−0.97, −0.44]; Table 3, Fig. 3A), indicating that chimpanzees were more likely to rest on the ground under warmer conditions (Fig. 3B). Rainfall, water vapor pressure, wind speed, and fruit availability showed no clear associations with substrate use. Previous-scan tree use was strongly associated with current tree use (β = 3.01, 95% CrI [2.54, 3.50]), indicating marked short-term temporal persistence in arboreal versus terrestrial resting. Day-level variation was also substantial (SD = 1.84, 95% CrI [1.32, 2.50]).

**Figure 3.**
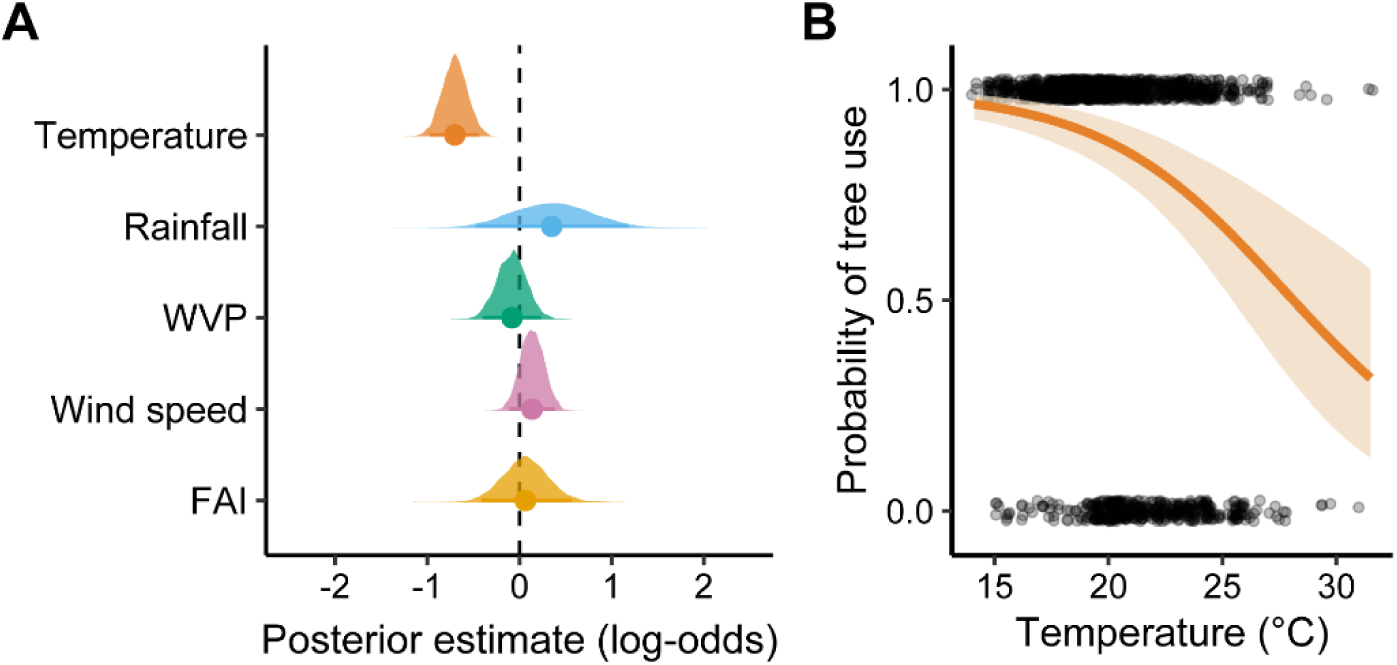
Effects of microclimatic variables on tree use during resting. (A) Posterior estimates from the Bayesian binomial GLMM for the effects of temperature, rainfall, water vapour pressure (WVP), wind speed, and fruit availability index (FAI) on tree use. Points indicate posterior means, shaded distributions show posterior uncertainty, and the dashed vertical line indicates no effect. Positive estimates indicate increased odds of tree use, whereas negative estimates indicate reduced odds. (B) Predicted probability of tree use across the observed temperature range. The line shows the posterior mean prediction and the shaded band represents the 95% credible interval. Points show observed individual records, with tree use coded as 1 and ground use as 0.

**Table 3.** Posterior estimates from a Bayesian binomial GLMM examining substrate use (tree vs. ground) by chimpanzees during resting in relation to microclimatic variables, fruit availability, and temporal dependence. Estimates are presented on the log-odds scale, with positive values indicating an increased probability of tree use relative to ground use. The lagged term (previous tree use) accounts for temporal dependence between consecutive observations. Day was included as a random intercept, with the random effect reported as the standard deviation (SD). Model convergence was satisfactory based on R^ values and effective sample sizes.

| Predictor | Estimate | Est. error | Lower 95% CrI | Upper 95% CrI | $\hat{R}$ |
| --- | --- | --- | --- | --- | --- |
| Intercept | -0.33 | 0.35 | -0.99 | 0.38 | 1.00 |
| Temperature | **-0.70** | 0.14 | **-0.97** | **-0.44** | 1.00 |
| Rainfall (yes) | 0.35 | 0.43 | -0.47 | 1.18 | 1.00 |
| Water vapour pressure | -0.09 | 0.16 | -0.40 | 0.23 | 1.00 |
| Wind speed | 0.14 | 0.13 | -0.11 | 0.39 | 1.00 |
| Fruit availability index | 0.07 | 0.25 | -0.41 | 0.57 | 1.00 |
| Previous tree use | **3.01** | 0.25 | **2.54** | **3.50** | 1.00 |
| **Random effect** |  |  |  |  |  |
| SD (day intercept) | 1.84 | 0.30 | 1.32 | 2.50 | 1.00 |

The canopy-strata results were most strongly associated with temperature (Table 4; Fig. 4A). Relative to the low canopy, the probabilities of using the mid- and top-canopy strata declined with increasing temperature (mid: β = −0.33, 95% CrI [−0.60, −0.08]; top: β = −0.59, [−0.94, −0.28]) (Fig. 4B). Wind speed was also negatively associated with top-canopy use (β = −0.31, 95% CrI [−0.61, −0.01]), whereas rainfall, WVP, and fruit availability showed no clear associations with canopy position. Previous-scan canopy composition indicated short-term temporal dependence. Mid-canopy use was more likely when a greater proportion of chimpanzees had occupied the mid (β = 1.73, 95% CrI [1.09, 2.38]) or top canopy (β = 1.30, [0.61, 2.04]) in the previous scan. Top-canopy use was likewise more likely following scans with higher proportions of chimpanzees in the mid (β = 0.85, [0.07, 1.62]) or top canopy (β = 2.57, [1.73, 3.42]). Day-level variation was substantial for both mid-(SD = 1.03, 95% CrI [0.67, 1.46]) and top-canopy use (SD = 1.49, [0.96, 2.16]).

**Figure 4.**
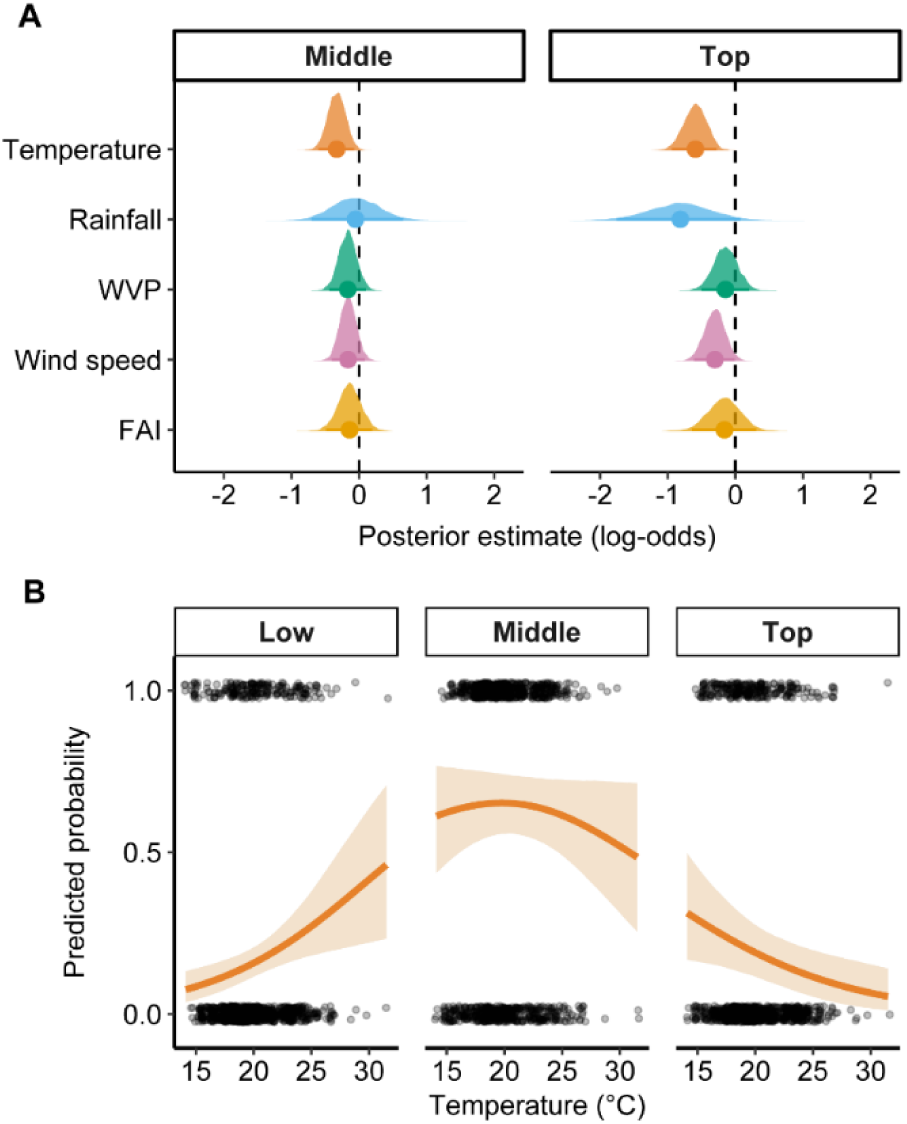
Effects of microclimatic variables on canopy-stratum use during arboreal resting. (A) Posterior estimates from the Bayesian multinomial GLMM for the effects of temperature, rainfall, water vapour pressure (WVP), wind speed, and fruit availability index (FAI) on use of the middle and top canopy relative to the low canopy. Points indicate posterior means, shaded distributions show posterior uncertainty, and the dashed vertical line indicates no effect. Positive estimates indicate increased odds of using that canopy stratum relative to the low canopy, whereas negative estimates indicate decreased odds. (B) Predicted probabilities of using the low, middle, and top canopy strata across the observed temperature range. Lines show posterior mean predictions and shaded bands represent 95% credible intervals. Points show observed individual canopy-stratum records.

**Table 4.** Posterior estimates from a Bayesian GLMM examining canopy strata use (low, mid, and top) by chimpanzees during resting in relation to microclimatic variables, fruit availability, and temporal dependence. Low canopy (L) was used as the reference category. Continuous predictors were standardized (mean = 0, SD = 1). Estimates are presented on the log-odds scale relative to low canopy. The lagged terms for previous canopy strata account for temporal dependence between consecutive observations. Day was included as a random intercept, with random effects reported as standard deviations (SD). Model convergence was satisfactory based on R^ values and effective sample sizes.

| Response category | Predictor | Estimate | Est. error | Lower 95% CrI | Upper 95% CrI | $\hat{R}$ |
| --- | --- | --- | --- | --- | --- | --- |
| **Mid** | Intercept | 0.09 | 0.30 | -0.49 | 0.71 | 1.00 |
|  | Temperature | **-0.33** | 0.13 | **-0.60** | **-0.08** | 1.00 |
|  | Rainfall (yes) | -0.05 | 0.35 | -0.70 | 0.65 | 1.00 |
|  | Water vapour pressure | -0.17 | 0.14 | -0.44 | 0.10 | 1.00 |
|  | Wind speed | -0.16 | 0.12 | -0.40 | 0.08 | 1.00 |
|  | Fruit availability index | -0.14 | 0.17 | -0.49 | 0.20 | 1.00 |
|  | Previous strata: mid | **1.73** | 0.33 | **1.09** | **2.38** | 1.00 |
|  | Previous strata: top | **1.30** | 0.37 | **0.61** | **2.04** | 1.00 |
| **Top** | Intercept | -1.00 | 0.39 | -1.78 | -0.24 | 1.00 |
|  | Temperature | **-0.59** | 0.17 | **-0.94** | **-0.28** | 1.00 |
|  | Rainfall (yes) | -0.82 | 0.47 | -1.76 | 0.07 | 1.00 |
|  | Water vapour pressure | -0.15 | 0.18 | -0.50 | 0.21 | 1.00 |
|  | Wind speed | **-0.31** | 0.15 | **-0.61** | **-0.01** | 1.00 |
|  | Fruit availability index | -0.16 | 0.24 | -0.64 | 0.30 | 1.00 |
|  | Previous strata: mid | **0.85** | 0.40 | **0.07** | **1.62** | 1.00 |
|  | Previous strata: top | **2.57** | 0.43 | **1.73** | **3.42** | 1.00 |
| **Random effects** | SD (day): mid strata | 1.03 | 0.20 | 0.67 | 1.46 | 1.00 |
|  | SD (day): top strata | 1.49 | 0.31 | 0.96 | 2.16 | 1.00 |

Day-nest use was strongly associated with temperature (Table 5; Fig. 5A). The probability of using a day nest declined as temperature increased (β = −0.43, 95% CrI [−0.71, −0.15]), indicating greater use of day nests under cooler conditions (Fig. 5B). In contrast, rainfall, fruit availability, water vapor pressure, and wind speed showed no clear associations with day-nest use, as their 95% credible intervals overlapped zero. Previous-scan day-nest use was also strongly associated with current use (β = 3.27, 95% CrI [2.78, 3.78]), indicating pronounced short-term temporal persistence in this behavior across successive scans. Day identity accounted for additional variation in day-nest use (SD = 1.55, 95% CrI [1.07, 2.15]), suggesting substantial between-day heterogeneity beyond the measured environmental predictors.

**Figure 5.**
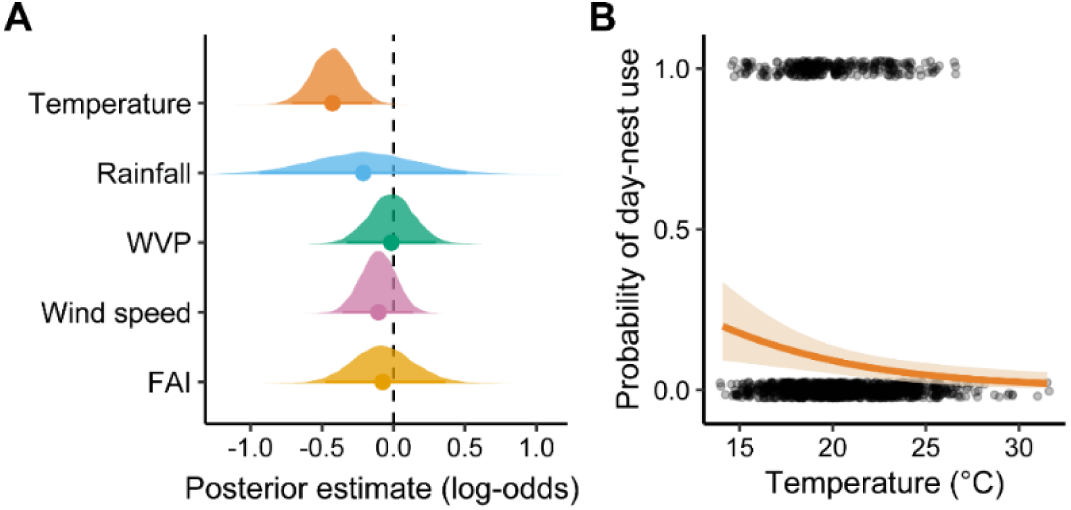
Effects of microclimatic variables on day-nest use during resting. (A) Posterior estimates from the Bayesian binomial GLMM for the effects of temperature, rainfall, water vapour pressure (WVP), wind speed, and fruit availability index (FAI) on day-nest use. Points indicate posterior means, shaded distributions show posterior uncertainty, and the dashed vertical line indicates no effect. Positive estimates indicate increased odds of day-nest use, whereas negative estimates indicate decreased odds. (B) Predicted probability of day-nest use across the observed temperature range. The line shows the posterior mean prediction and the shaded band represents the 95% credible interval. Points show observed individual records, with day-nest use coded as 1 and non-use as 0.

**Table 5.** Posterior estimates from a Bayesian binomial GLMM examining day-nest use by chimpanzees during resting in relation to microclimatic variables, fruit availability, and previous day-nest use. Day-nest use was coded as 1 and non-nest resting as 0. Continuous predictors were standardized before analysis. Estimates are presented on the log-odds scale with 95% credible intervals (CrIs). The lagged term (previous day-nest use) accounts for temporal dependence between consecutive observations.

| Predictor | Estimate | Est. error | Lower 95% CrI | Upper 95% CrI | $\hat{R}$ |
| --- | --- | --- | --- | --- | --- |
| Intercept | -3.10 | 0.28 | -3.69 | -2.59 | 1.00 |
| Temperature | **-0.43** | 0.14 | **-0.71** | **-0.15** | 1.00 |
| Rainfall (yes) | -0.21 | 0.37 | -0.94 | 0.51 | 1.00 |
| Fruit availability index | -0.07 | 0.21 | -0.48 | 0.36 | 1.00 |
| Water vapour pressure | -0.02 | 0.16 | -0.33 | 0.30 | 1.00 |
| Wind speed | -0.11 | 0.13 | -0.36 | 0.14 | 1.00 |
| Previous day-nest use | **3.27** | 0.25 | **2.78** | **3.78** | 1.00 |
| **Random effect** |  |  |  |  |  |
| SD (day intercept) | 1.55 | 0.28 | 1.07 | 2.15 | 1.00 |

## 4 Discussion

Our results show that temperature was a key factor that shaped how Nyungwe chimpanzees rest. Across several behaviors, chimpanzees changed their posture, rest site, height in the tree canopy, and use of a day nest in ways that are consistent with behavioral thermoregulation. In cooler conditions, they were more likely to curl, huddle, or rest in a day nest. In warmer conditions, they were more likely to rest in shade, rest on the ground, and avoid the higher canopy. These results support the idea that animals can choose their rest sites in a way that helps them to conserve heat when it is cool and reduce heat gain when it is warm. Rest in this montane population, therefore, does not appear to be simply a period of inactivity. Instead, it may provide an important context in which chimpanzees adjust body posture, sun exposure, substrate use, and their choice of height in a tree in response to local thermal conditions.

Of all the measured weather variables, resting posture was associated mainly with ambient temperature and, to a lesser extent, water vapor pressure (WVP), whereas rainfall, wind speed, and fruit availability showed little clear effect. Chimpanzees were more likely to adopt curled and huddled postures under cooler conditions, and the probabilities of these postures declined as temperature increased. This pattern is consistent with behavioral thermoregulation because primates can reduce heat loss in cold conditions by adopting compact postures, increasing body contact, and reducing exposed body surface area (Stelzner & Hausfater, 1986; Terrien et al., 2011; Thompson & Hermann, 2024). Similar responses have been reported in other primates. Yellow baboons in Amboseli National Park use hunched postures during cool mornings (Stelzner & Hausfater, 1986), while collared brown lemurs curl more frequently during cooler periods and ring-tailed lemurs use sunning behavior during cold mornings and winter months (Donati et al., 2011; Kelley et al., 2016). Temperature-related huddling has also been reported in Barbary macaques (Campbell et al., 2018b), red-fronted lemurs (Ostner, 2002), and Japanese macaques (Ogawa & Wada, 2011; Hanya et al., 2007; Ishizuka, 2021). Huddling was also less likely under conditions of higher WVP. In Nyungwe’s cool and humid montane forest, higher atmospheric moisture may reduce evaporative heat loss, as high humidity can constrain evaporative cooling in primates (Friedl & Holmes, 1986). Because huddling reduces exposed surface area and promotes heat conservation, close body contact may therefore become less thermally advantageous under more humid conditions. However, this interpretation remains tentative, particularly because huddling may also reflect social integration, affiliation, and partner choice rather than thermoregulation alone (Gestich et al., 2014; Kelley et al., 2016; McFarland et al., 2015). Lying also became less likely relative to sitting as temperature increased, although its association with temperature was weaker than those of curled and huddled postures. This suggests that lying may also be influenced by thermal conditions, but that curling and huddling represent stronger responses to cooler temperatures. Although we expected wind speed to influence postural behavior, neither wind speed nor rainfall showed clear associations with posture. This does not mean that these variables are unimportant for postural thermoregulation more broadly. For example, yellow baboons orient their furry backs toward cool air and reduce exposure of the less-furred ventrum and face (Stelzner & Hausfater, 1986). Overall, therefore, temperature was the dominant climatic correlate of resting posture in Nyungwe chimpanzees, with WVP showing an additional, more specific association with huddling.

The microhabitat used during rest also changed with temperature. As temperature increased, chimpanzees were more likely to rest in shade and less likely to rest in trees, an indication that they adjusted both sun exposure and substrate use under warmer conditions. More use of shade is consistent with heat avoidance because shade reduces direct solar radiation and radiant heat load, and helps animals to limit heat gain during warm periods (Hill et al., 2004; Norris & Kunz, 2012). Similar sun/shade responses have been reported in other primates. Yellow baboons avoid direct sun and use shade during hot conditions but increase sun exposure when conditions are cooler (Stelzner & Hausfater, 1986). Tibetan macaques avoid direct sunlight and rest in dense forest during hot summer periods but bask on exposed rocks during cold winter months (Zhou et al., 2022). Black-fronted titi monkeys also spend more time in sunny places during cooler conditions and shift towards shaded sites as temperatures increase (Gestich et al., 2014). Although the effect of humidity on sun/shade use did not support our hypothesis, sun-exposure behavior in some primates is shaped not only by temperature but also by humidity or seasonal context. For example, humidity influenced sun avoidance in yellow baboons, particularly during the lush season, whereas temperature was more important during the dry season (Pochron, 2000). Similarly, Japanese macaques selected shade under hot and humid conditions but used semi-shade under hot and dry conditions (Tabuse, 2026). In the Nyungwe chimpanzees, by contrast, sun/shade selection showed a clearer association with temperature but not with humidity or wind. This suggests that, within the range of conditions that we observed in this montane forest, temperature was the main climatic cue that shaped sun-exposure decisions during rest.

The chimpanzees used trees for resting less under warmer conditions than under cooler conditions, which suggests that ground resting may form part of a broader heat-avoidance response, consistent with the observed increase in shade use. Thermal conditions vary vertically within forests, and primates can experience different levels of solar exposure, wind, humidity, and heat at different canopy heights (Thompson et al., 2016). Comparative work across arboreal primates shows that higher temperatures are often associated with more ground use, suggesting that terrestriality may help some species to cope with warmer or more exposed conditions (Eppley et al., 2022). Chimpanzee studies support this interpretation: in Budongo Forest, chimpanzees reduced prolonged exposure to direct sun and increased ground use as temperature rose (Kosheleff & Anderson, 2009), while at Bossou, chimpanzees spent more time on the ground during warm, dry months, a pattern linked to local forest microclimate (Takemoto, 2004). In Nyungwe, greater ground-based rest under warmer conditions may therefore allow chimpanzees to use cooler, more shaded, or less exposed sites for rest, although this interpretation remains tentative because we did not directly measure thermal differences between the canopy and the ground. Although temperature was the main predictor of tree– ground rest in our study, food availability and rainfall may still influence the use of vertical space indirectly or under different ecological conditions. Previous studies suggest that primates may increase ground use when arboreal food resources are scarce: ground use in titi monkeys has been linked to seasonal reductions in arboreal fruit availability, and terrestrial feeding in Coimbra-Filho’s titi monkeys was associated with the absence of arboreal fruit and young leaf resources (Souza-Alves et al., 2019, 2021). Comparative evidence also suggests that species that rely less on fruit tend to show greater ground use, indicating that reduced fruit availability or lower frugivory may promote terrestriality in some contexts (Eppley et al., 2022). Rainfall may also affect tree–ground use indirectly. In titi monkeys, lower rainfall, used as a proxy for stronger seasonality and reduced fruit availability, was associated with more frequent terrestrial behaviour (Souza-Alves et al., 2019). Dry-season or low-water conditions can also lead arboreal primates to descend to the ground to access terrestrial water sources, as reported for northern muriquis, Temminck’s red colobus, buffy-headed marmosets, and red-fronted lemurs (Mourthé et al., 2007; Ferrari & Hilário, 2012; Hillyer et al., 2015; Amoroso et al., 2020).

Whether chimpanzees adjust their height within the canopy during rest has not been examined. To our knowledge, no previous study has directly tested whether chimpanzees choose different canopy strata during rest in relation to measured microclimatic conditions. In Nyungwe, canopy-stratum use varied with temperature: as temperature increased, chimpanzees were less likely to rest in the middle or upper canopy and more likely to rest in the lower canopy. This pattern is consistent with the observed increase in ground use under warmer conditions and suggests that a downward shift may form part of a broader heat-avoidance response. By moving away from higher levels of the canopy, chimpanzees may reduce exposure to direct solar radiation and use cooler, more shaded, or more thermally buffered parts of the forest (Thompson et al., 2016). Evidence from other arboreal primates suggests that vertical position can contribute to thermoregulatory microhabitat selection, although the direction of response may vary among species and habitats. For example, black-fronted titi monkeys adjusted forest-stratum use in relation to thermal conditions and solar exposure during inactive periods (Gestich et al., 2014), while mantled howler monkeys combined postural adjustment with changes in tree-stratum use and selected lower levels for rest under warmer conditions and higher levels under cooler conditions (Palestino-Sánchez et al., 2025b). However, because direct evidence for thermoregulatory use of canopy strata in wild chimpanzees is lacking, the downward shift observed in Nyungwe should be interpreted as a possible thermoregulatory response rather than definitive evidence. Wind speed was also negatively associated with use of the upper canopy, suggesting that chimpanzees may avoid more exposed canopy positions under windier conditions. This pattern is consistent with the prediction that lower canopy strata may provide greater shelter from convective heat loss. Rainfall, by contrast, showed no clear association with canopy-stratum use. Given that observations in the upper canopy were limited, we cannot interpret the wind speed effect as strong evidence of avoidance of the upper canopy. Further work that directly measures wind exposure and thermal conditions across canopy strata would help determine whether chimpanzees systematically shift downward to reduce exposure under windy conditions.

The use of a day nest during rest was also associated with temperature, suggesting that nests may provide a thermally beneficial structure for rest under cooler conditions. Chimpanzees were more likely to use a day nest when temperature was low, and day-nest use declined as temperature increased. This pattern is consistent with the idea that great ape nests provide more than support or comfort and may also help buffer individuals from unfavorable thermal conditions (Fruth & Hohmann, 1996; Anderson, 2000; Stewart, 2011). Most evidence for the thermal function of ape nests comes from studies of night nests, including work showing that chimpanzees adjust nest structure in response to local weather (Stewart et al., 2018; Al-Razi et al., 2026). Much less is known about the function of day nests, which are usually used for shorter periods of rest and may serve several functions, including comfort, support, and thermoregulation (van Lawick-Goodall, 1968; McGrew, 2021). Therefore, the increased use of day nests under cooler conditions in Nyungwe should be interpreted as suggestive rather than definitive evidence that they provide thermal buffering. Rainfall showed no clear association with day-nest use, providing little evidence that day nests were used specifically in response to daytime rainfall. Future work should examine day-nest structure, the microclimate around the nest, canopy cover, and body posture during day-nest use to test more directly whether these structures provide thermal benefits during daytime rest.

## 5 Conclusions

Overall, of the variables that we measured, temperature-based predictions received the strongest and most consistent support, whereas the effects of wind speed, WVP, rainfall, and fruit availability were more limited or context-dependent. This pattern may indicate that, during rest, chimpanzees respond most directly to ambient temperature, while other climatic variables influence more specific aspects of posture or microhabitat use. For example, higher WVP was associated with reduced huddling, while higher wind speed was associated with reduced use of the upper canopy. Wind and rainfall can also be buffered by canopy cover, vegetation density, slope, and the precise position of the resting individual, making their effects more difficult to detect when using scan-level weather measurements. Similarly, fruit availability may shape ranging and feeding opportunities more than the immediate resting posture or the choice of resting site. The generally limited or behavior-specific effects of wind, WVP, rainfall, and fruit availability do not imply that these variables are unimportant, but rather that their influence may be indirect, context-dependent, or expressed only in particular aspects of resting behavior. Although the observed patterns are consistent with behavioral thermoregulation, the decisions about rest may also be shaped by habitat structure, fruit availability, predation risk, social context, and previous activity (Isbell & Young, 1993; Di Bitetti et al., 2000; Korstjens et al., 2010; Bryson-Morrison et al., 2017; Eppley et al., 2022). Future work that combines behavioral observations with direct microclimate measurements at the resting site, habitat structure, fruit and non-fruit food availability, social context, and previous activity would help clarify the relative importance of thermal and non-thermal mechanisms in the chimpanzee resting behavior.

## Author Contributions

**Hassan Al-Razi:** conceptualization, methodology, investigation, data curation, formal analysis, software, visualization, funding acquisition, writing – original draft, writing – review and editing. **Fidele Muhayeyezu:** data collection, project administration. **Felix Mulindahabi:** project administration, writing – review and editing. **Protais Niyigaba:** project administration, writing – review and editing. **Drew Arthur Bantlin:** project administration, writing – review and editing. **Beth A. Kaplin:** methodology, supervision, writing – original draft, writing – review and editing. **Shane K. Maloney:** administration, supervision, writing – original draft, writing – review and editing. **Cyril C. Grueter:** conceptualization, methodology, supervision, funding acquisition, writing – original draft, writing – review and editing.

## Acknowledgements

We thank Dr. Richard Muvunyi and Albert Kayitare at the Rwanda Development Board (RDB) for permission to conduct research in Nyungwe National Park. We are grateful to the African Parks trackers for their assistance in locating chimpanzees, and to Peter Niyonzima and Philbert Muhire for their support with chimpanzee tracking and data collection. We also thank African Parks for providing accommodation at Mayebe camp. All research was conducted in accordance with the legal requirements of Rwanda and RDB regulations. Funding was provided by The University of Western Australia, the Primate Action Fund, the Primate Society of Great Britain, Basler Stiftung für biologische Forschung, and Stiftung Temperatio.

## Data Availability Statement

The data that support the findings of this study are available in the supporting material of this article.

## References

Al-Razi, H., F. Muhayeyezu, F. Mulindahabi, P. Niyigaba, D. A. Bantlin, L. Ding, B. A. Kaplin, S. K. Maloney, and C. C. Grueter. 2026. “Thermal Adaptation and the Potential Anticipation of Overnight Weather in the Nesting Decisions of Chimpanzees.” Current Biology 36: 2662–2672.e5. 10.1016/j.cub.2026.04.005.

Altmann, J. 1974. “Observational Study of Behavior: Sampling Methods.” Behaviour 49, no. 3/4: 227–265. 10.1163/156853974X00534.

Amoroso, C. R., P. M. Kappeler, C. Fichtel, and C. L. Nunn. 2020. “Water Availability Impacts Habitat Use by Red-Fronted Lemurs (*Eulemur rufifrons*): An Experimental and Observational Study.” International Journal of Primatology 41: 61–80. 10.1007/s10764-020-00136-9.

Anderson, J. R. 2000. “Sleep-Related Behavioural Adaptations in Free-Ranging Anthropoid Primates.” Sleep Medicine Reviews 4, no. 4: 355–373. 10.1053/smrv.2000.0105.

Barenbrug, A. W. T. 1974. Psychrometry and Psychrometric Charts. 3rd ed. Johannesburg: Chamber of Mines of South Africa.

Bateson, M., and P. Martin. 2021. Measuring Behaviour: An Introductory Guide. 4th ed. Cambridge: Cambridge University Press.

Bicca-Marques, J. C., and C. Calegaro-Marques. 1998. “Behavioral Thermoregulation in a Sexually and Developmentally Dichromatic Neotropical Primate, the Black-and-Gold Howling Monkey (*Alouatta caraya*).” American Journal of Physical Anthropology 106: 533–546. 10.1002/(SICI)1096-8644(199808)106:4<533::AID-AJPA8>3.0.CO;2-J.

Boesch, C. 1995. “Innovation in Wild Chimpanzees (*Pan troglodytes*).” International Journal of Primatology 16: 1–15. 10.1007/BF02700150.

Brividoro, M. V., L. I. Oklander, V. I. Cantarelli, M. F. Ponzio, H. R. Ferrari, and M. M. Kowalewski. 2023. “Influence of Weather Conditions on Sleeping Patterns and Selection of Foliage Cover of Sleeping Trees in Black-and-Gold Howler Monkeys (*Alouatta caraya*) in Northern Argentina.” International Journal of Primatology 44, no. 6: 1110–1126.

Brown, M., and C. T. Downs. 2002. “The Role of Shading Behavior in the Thermoregulation of Breeding Crowned Plovers (*Vanellus coronatus*).” Journal of Thermal Biology 28: 51–58.

Brownlow, A. R., A. J. Plumptre, V. Reynolds, and R. Ward. 2001. “Sources of Variation in the Nesting Behavior of Chimpanzees (*Pan troglodytes schweinfurthii*) in the Budongo Forest, Uganda.” American Journal of Primatology 55: 49–55. 10.1002/ajp.1038.

Bruijnzeel, L. A., F. N. Scatena, and L. S. Hamilton, eds. 2010. Tropical Montane Cloud Forests: Science for Conservation and Management. Cambridge: Cambridge University Press.

Bryson-Morrison, N., J. Tzanopoulos, T. Matsuzawa, and T. Humle. 2017. “Activity and Habitat Use of Chimpanzees (*Pan troglodytes verus*) in the Anthropogenic Landscape of Bossou, Guinea, West Africa.” International Journal of Primatology 38: 282–302. 10.1007/s10764-016-9947-4.

Bürkner, P.-C. 2017. “brms: An R Package for Bayesian Multilevel Models Using Stan.” Journal of Statistical Software 80: 1–28. 10.18637/jss.v080.i01.

Campbell, L. A. D., P. J. Tkaczynski, M. Mouna, A. Derrou, L. Oukannou, B. Majolo, and E. Van Lavieren. 2018a. “Behavioural Thermoregulation via Microhabitat Selection of Winter Sleeping Areas in an Endangered Primate: Implications for Habitat Conservation.” Royal Society Open Science 5: 181113. 10.1098/rsos.181113.

Campbell, L. A. D., P. J. Tkaczynski, J. Lehmann, M. Mouna, and B. Majolo. 2018b. “Social Thermoregulation as a Potential Mechanism Linking Sociality and Fitness: Barbary Macaques With More Social Partners Form Larger Huddles.” Scientific Reports 8: 6074. 10.1038/s41598-018-24373-4.

Carrascal, L. M., J. A. Díaz, D. L. Huertas, and I. Mozetich. 2001. “Behavioral Thermoregulation by Treecreepers: Trade-Off Between Saving Energy and Reducing Crypsis.” Ecology 82: 1642–1654.

Chapman, C. A., L. J. Chapman, and R. W. Wrangham. 1995. “Ecological Constraints on Group Size: An Analysis of Spider Monkey and Chimpanzee Subgroups.” Behavioral Ecology and Sociobiology 36: 59–70. 10.1007/BF00175729.

Chapman, H., N. J. Cordeiro, P. Dutton, D. Wenny, S. Kitamura, B. Kaplin, F. P. L. Melo, and M. J. Lawes. 2016. “Seed-Dispersal Ecology of Tropical Montane Forests.” Journal of Tropical Ecology 32: 437–454. 10.1017/S0266467416000389.

Chen-Kraus, C., N. A. Raharinoro, R. R. Lawler, and A. F. Richard. 2023. “Terrestrial Tree Hugging in a Primarily Arboreal Lemur (*Propithecus verreauxi*): A Cool Way to Deal With Heat?” International Journal of Primatology 44, no. 1: 178–191.

DaSilva, G. L. 1993. “Postural Changes and Behavioural Thermoregulation in Colobus polykomos: The Effect of Climate and Diet.” African Journal of Ecology 31: 226–241. 10.1111/j.1365-2028.1993.tb00536.x.

Di Bitetti, M. S., E. M. Luengos Vidal, M. C. Baldovino, and V. Benesovsky. 2000. “Sleeping Site Preferences in Tufted Capuchin Monkeys (*Cebus apella nigritus*).” American Journal of Primatology 50, no. 4: 257–274. 10.1002/(SICI)1098-2345(200004)50:4<257::AID-AJP3>3.0.CO;2-J.

Donati, G., E. Ricci, N. Baldi, V. Morelli, and S. M. Borgognini-Tarli. 2011. “Behavioral Thermoregulation in a Gregarious Lemur, Eulemur collaris: Effects of Climatic and Dietary-Related Factors.” American Journal of Physical Anthropology 144: 355–364. 10.1002/ajpa.21415.

Dugger, P. J., N. J. Cordeiro, F. Mulindahabi, M. Bana, and B. A. Kaplin. 2025. “Increasing Fruiting Synchrony at the Community Level in an Afromontane Tropical Forest.” Ecosphere 16: e70409. 10.1002/ecs2.70409.

Duncan, L. M., and N. Pillay. 2013. “Shade as a Thermoregulatory Resource for Captive Chimpanzees.” Journal of Thermal Biology 38, no. 4: 169–177. 10.1016/j.jtherbio.2013.02.009.

Eppley, T. M., S. Hoeks, C. A. Chapman, J. U. Ganzhorn, K. Hall, M. A. Owen, D. B. Adams, et al. 2022. “Factors Influencing Terrestriality in Primates of the Americas and Madagascar.” Proceedings of the National Academy of Sciences 119: e2121105119. 10.1073/pnas.2121105119.

Fashing, P. J. 2001. “Activity and Ranging Patterns of Guerezas in the Kakamega Forest: Intergroup Variation and Implications for Intragroup Feeding Competition.” International Journal of Primatology 22: 549–577.

Ferrari, S. F., and R. R. Hilário. 2012. “Use of Water Sources by Buffy-Headed Marmosets (*Callithrix flaviceps*) at Two Sites in the Brazilian Atlantic Forest.” Primates 53: 65–70. 10.1007/s10329-011-0277-z.

Fox, J., and S. Weisberg. 2019. An R Companion to Applied Regression. 3rd ed. Thousand Oaks, CA: Sage.

Friedl, K. E., and W. N. Holmes. 1986. “The Effect of Relative Humidity on Osmoregulation in the Squirrel Monkey (*Saimiri sciureus*).” Primates 27, no. 4: 465–470. 10.1007/BF02381891.

Fruth, B., and G. Hohmann. 1996. “Nest Building Behavior in the Great Apes: The Great Leap Forward?” In Great Ape Societies, edited by W. C. McGrew, L. F. Marchant, and T. Nishida, 225–240. Cambridge: Cambridge University Press.

Fruth, B., and W. C. McGrew. 1998. “Resting and Nesting in Primates: Behavioral Ecology of Inactivity.” American Journal of Primatology 46: 3–5.

Gestich, C. C., C. B. Caselli, and E. Z. F. Setz. 2014. “Behavioural Thermoregulation in a Small Neotropical Primate.” Ethology 120: 331–339. 10.1111/eth.12203.

Gilby, I. C., A. A. Pokempner, and R. W. Wrangham. 2010. “A Direct Comparison of Scan and Focal Sampling Methods for Measuring Wild Chimpanzee Feeding Behaviour.” Folia Primatologica 81, no. 5: 254–264. 10.1159/000322354.

Green, S. J., B. J. Boruff, and C. C. Grueter. 2020a. “From Ridgetops to Ravines: Landscape Drivers of Chimpanzee Ranging Patterns.” Animal Behaviour 163: 51–60. 10.1016/j.anbehav.2020.02.016.

Green, S. J., B. J. Boruff, T. R. Bonnell, and C. C. Grueter. 2020b. “Chimpanzees Use Least-Cost Routes to Out-of-Sight Goals.” Current Biology 30: 4528–4533.e5. 10.1016/j.cub.2020.08.076.

Gross-Camp, N., and B. A. Kaplin. 2005. “Chimpanzee (*Pan troglodytes*) Seed Dispersal in an Afromontane Forest: Microhabitat Influences on the Postdispersal Fate of Large Seeds.” Biotropica 37: 641–649. 10.1111/j.1744-7429.2005.00081.x.

Grueter, C. C., D. Li, B. Ren, and M. Li. 2013. “Substrate Use and Postural Behavior in Free-Ranging Snub-Nosed Monkeys (*Rhinopithecus bieti*) in Yunnan.” Integrative Zoology 8: 335–345. 10.1111/1749-4877.12023.

Hanya, G., M. Kiyono, and S. Hayaishi. 2007. “Behavioral Thermoregulation of Wild Japanese Macaques: Comparisons Between Two Subpopulations.” American Journal of Primatology 69: 802–815. 10.1002/ajp.20397.

Henzi, S. P., R. Hetem, A. Fuller, S. Maloney, C. Young, D. Mitchell, L. Barrett, and R. McFarland. 2017. “Consequences of Sex-Specific Sociability for Thermoregulation in Male Vervet Monkeys During Winter.” Journal of Zoology 302: 193–200. 10.1111/jzo.12448.

Herbers, J. M. 1981. “Time Resources and Laziness in Animals.” Oecologia 49: 252–262. 10.1007/BF00349198.

Hill, R. A. 1999. “Ecological and Demographic Determinants of Time Budgets in Baboons: Implications for Cross-Populational Models of Baboon Socioecology.” PhD diss., University of Liverpool.

Hill, R. A., T. Weingrill, L. Barrett, and S. P. Henzi. 2004. “Indices of Environmental Temperatures for Primates in Open Habitats.” Primates 45: 7–13.

Hillyer, A. P., R. Armstrong, and A. H. Korstjens. 2015. “Dry Season Drinking From Terrestrial Man-Made Watering Holes in Arboreal Wild Temminck’s Red Colobus, The Gambia.” Primate Biology 2: 21–24. 10.5194/pb-2-21-2015.

Isbell, L. A., and T. P. Young. 1993. “Social and Ecological Influences on Activity Budgets of Vervet Monkeys, and Their Implications for Group Living.” Behavioral Ecology and Sociobiology 32: 377–385. 10.1007/BF00168821.

Ishizuka, S. 2021. “Do Dominant Monkeys Gain More Warmth? Number of Physical Contacts and Spatial Positions in Huddles for Male Japanese Macaques in Relation to Dominance Rank.” Behavioural Processes 185: 104317. 10.1016/j.beproc.2021.104317.

Kelley, E. A., N. G. Jablonski, G. Chaplin, R. W. Sussman, and J. M. Kamilar. 2016. “Behavioral Thermoregulation in *Lemur catta*: The Significance of Sunning and Huddling Behaviors.” American Journal of Primatology 78: 745–754. 10.1002/ajp.22538.

Kluiver, C. E., J. J. M. Massen, and D. Bhattacharjee. 2022. “Personality as a Predictor of Time-Activity Budget in Lion-Tailed Macaques (*Macaca silenus*).” Animals 12, no. 12: 1495. 10.3390/ani12121495.

Koops, K., W. C. McGrew, H. de Vries, and T. Matsuzawa. 2012. “Nest-Building by Chimpanzees (*Pan troglodytes verus*) at Seringbara, Nimba Mountains: Antipredation, Thermoregulation, and Antivector Hypotheses.” International Journal of Primatology 33: 356–380. 10.1007/s10764-012-9585-4.

Korstjens, A. H., J. Lehmann, and R. I. M. Dunbar. 2010. “Resting Time as an Ecological Constraint on Primate Biogeography.” Animal Behaviour 79: 361–374. 10.1016/j.anbehav.2009.11.012.

Kosheleff, V. P., and C. N. K. Anderson. 2009. “Temperature’s Influence on the Activity Budget, Terrestriality, and Sun Exposure of Chimpanzees in the Budongo Forest, Uganda.” American Journal of Physical Anthropology 139: 172–181. 10.1002/ajpa.20970.

Li, D., B. Ren, C. C. Grueter, B. Li, and M. Li. 2010. “Nocturnal Sleeping Habits of the Yunnan Snub-Nosed Monkey in Xiangguqing, China.” American Journal of Primatology 72, no. 12: 1092–1099.

Li, Y. 2007. “Terrestriality and Tree Stratum Use in a Group of Sichuan Snub-Nosed Monkeys.” Primates 48: 197–207. 10.1007/s10329-006-0035-9.

Lopes, K. G., and J. C. Bicca-Marques. 2017. “Ambient Temperature and Humidity Modulate the Behavioural Thermoregulation of a Small Arboreal Mammal (*Callicebus bernhardi*).” Journal of Thermal Biology 69: 104–109. 10.1016/j.jtherbio.2017.06.010.

Lüdecke, D., M. S. Ben-Shachar, I. Patil, P. Waggoner, and D. Makowski. 2021. “performance: An R Package for Assessment, Comparison and Testing of Statistical Models.” Journal of Open Source Software 6, no. 60: 3139. 10.21105/joss.03139.

Matsuda, I., C. A. Chapman, C. Y. Shi Physilia, J. C. Mun Sha, and M. Clauss. 2017. “Primate Resting Postures: Constraints by Foregut Fermentation?” Physiological and Biochemical Zoology 90: 383–391. 10.1086/691360.

Matthews, J. K., A. Ridley, B. A. Kaplin, and C. C. Grueter. 2021. “Ecological and Reproductive Drivers of Fission-Fusion Dynamics in Chimpanzees (*Pan troglodytes schweinfurthii*) Inhabiting a Montane Forest.” Behavioral Ecology and Sociobiology 75, no. 1: 1–9.

Matthews, J. K., A. Ridley, P. Niyigaba, B. A. Kaplin, and C. C. Grueter. 2019. “Chimpanzee Feeding Ecology and Fallback Food Use in the Montane Forest of Nyungwe National Park, Rwanda.” American Journal of Primatology 81: e22971. 10.1002/ajp.22971.

McFarland, R., S. P. Henzi, L. Barrett, A. Wanigaratne, E. Coetzee, A. Fuller, R. S. Hetem, D. Mitchell, and S. K. Maloney. 2016. “Thermal Consequences of Increased Pelt Loft Infer an Additional Utilitarian Function for Grooming.” American Journal of Primatology 78: 456–461.

McFarland, R., L. Barrett, M. Costello, A. Fuller, R. S. Hetem, S. K. Maloney, D. Mitchell, and P. S. Henzi. 2020. “Keeping Cool in the Heat: Behavioral Thermoregulation and Body Temperature Patterns in Wild Vervet Monkeys.” American Journal of Physical Anthropology 171: 407–418. 10.1002/ajpa.23962.

McFarland, R., A. Fuller, R. S. Hetem, D. Mitchell, S. K. Maloney, S. P. Henzi, and L. Barrett. 2015. “Social Integration Confers Thermal Benefits in a Gregarious Primate.” Journal of Animal Ecology 84: 871–878. 10.1111/1365-2656.12329.

McGrew, W. C. 2021. “Sheltering Chimpanzees.” Primates 62: 445–455. 10.1007/s10329-021-00903-z.

Mekonnen, A., P. J. Fashing, E. J. Sargis, V. V. Venkataraman, A. Bekele, R. A. Hernandez-Aguilar, E. K. Rueness, and N. C. Stenseth. 2018. “Flexibility in Positional Behavior, Strata Use, and Substrate Utilization Among Bale Monkeys (*Chlorocebus djamdjamensis*) in Response to Habitat Fragmentation and Degradation.” American Journal of Primatology 80: e22760. 10.1002/ajp.22760.

Mourthé, Í. M. C., D. Guedes, J. Fidelis, J. P. Boubli, S. L. Mendes, and K. B. Strier. 2007. “Ground Use by Northern Muriquis (*Brachyteles hypoxanthus*).” American Journal of Primatology 69: 706–712. 10.1002/ajp.20405.

Norris, A. L., and T. H. Kunz. 2012. “Effects of Solar Radiation on Animal Thermoregulation.” In Solar Radiation, edited by E. B. Babatunde. IntechOpen. 10.5772/34771.

Nyirambangutse, B., E. Zibera, F. K. Uwizeye, D. Nsabimana, E. Bizuru, H. Pleijel, J. Uddling, and G. Wallin. 2017. “Carbon Stocks and Dynamics at Different Successional Stages in an Afromontane Tropical Forest.” Biogeosciences 14: 1285–1303. 10.5194/bg-14-1285-2017.

Ogawa, H., and K. Wada. 2011. “Shape of, and Body Direction in, Huddles of Japanese Macaques (*Macaca fuscata*) in Arashiyama, Japan.” Primates 52: 229–235. 10.1007/s10329-011-0248-4.

Ostner, J. 2002. “Social Thermoregulation in Redfronted Lemurs (*Eulemur fulvus rufus*).” Folia Primatologica 73: 175–180.

Palestino-Sánchez, R. M., N. S. Huerta-Pacheco, F. García-Orduña, J. F. Rodríguez-Landa, M. A. Cruz-Aguilar, and M. d. J. Rovirosa-Hernández. 2025a. “Effects of Fluctuation in Temperature and Humidity and Behavioral Thermoregulation on the Resting Time of *Alouatta palliata* Monkeys.” Animal Biodiversity and Conservation 48: e0008. 10.32800/abc.2025a.48.0008.

Palestino-Sánchez, R. M., N. S. Huerta-Pacheco, J. F. Rodríguez-Landa, M. F. López-Flores, F. García-Orduña, and M. d. J. Rovirosa-Hernández. 2025b. “Howler Monkey (*Alouatta palliata*) Thermal Behavioral Strategies and Resting Time Are Temperature-Dependent.” Current Zoology. 10.1093/cz/zoaf068.

Paterson, J. D. 1981. “Postural-Positional Thermoregulatory Behaviour and Ecological Factors in Primates.” Canadian Review of Physical Anthropology 3: 3–11.

Plumptre, A. J., M. Masozera, P. J. Fashing, A. McNeilage, C. Ewango, B. A. Kaplin, and I. Liengola. 2002. Biodiversity Surveys of the Nyungwe Forest Reserve in S.W. Rwanda. Wildlife Conservation Society Working Paper 19: 1–96.

Plumptre, A. J., R. Rose, G. Nagnendo, E. A. Williamson, K. Didier, J. Hart, F. Mulindahabi, C. Hicks, B. Griffin, H. Ogawa, et al. 2010. Eastern Chimpanzee (Pan troglodytes schweinfurthii): Status Survey and Conservation Action Plan 2010–2020. Gland, Switzerland: IUCN.

Pochron, S. T. 2000. “Sun Avoidance in the Yellow Baboons (*Papio cynocephalus cynocephalus*) of Ruaha National Park, Tanzania: Variations With Season, Behavior and Weather.” International Journal of Biometeorology 44: 141–147. 10.1007/s004840000058.

Pruetz, J. D. 2007. “Evidence of Cave Use by Savanna Chimpanzees (*Pan troglodytes verus*) at Fongoli, Senegal: Implications for Thermoregulatory Behavior.” Primates 48: 316–319. 10.1007/s10329-007-0038-1.

R Core Team. 2023. R: A Language and Environment for Statistical Computing. Vienna: R Foundation for Statistical Computing. https://www.R-project.org/.

Roberts, S. C., and R. I. M. Dunbar. 1991. “Climatic Influences on the Behavioural Ecology of Chanter’s Mountain Reedbuck in Kenya.” African Journal of Ecology 29: 316–329. 10.1111/j.1365-2028.1991.tb00469.x.

Samson, D. R., and K. D. Hunt. 2012. “A Thermodynamic Comparison of Arboreal and Terrestrial Sleeping Sites for Dry-Habitat Chimpanzees (*Pan troglodytes schweinfurthii*) at the Toro-Semliki Wildlife Reserve, Uganda.” American Journal of Primatology 74: 811–818. 10.1002/ajp.22031.

Sharma, N., M. A. Huffman, S. Gupta, H. Nautiyal, R. Mendonça, L. Morino, and A. Sinha. 2016. “Watering Holes: The Use of Arboreal Sources of Drinking Water by Old World Monkeys and Apes.” Behavioural Processes 129: 18–26. 10.1016/j.beproc.2016.05.006

Shukla, I., A. M. Kilpatrick, and R. S. Beltran. 2021. “Variation in Resting Strategies Across Trophic Levels and Habitats in Mammals.” Ecology and Evolution 11: 14405–14415. 10.1002/ece3.8073.

Smith, M. H. 1983. “Strong Selection and Rapid Evolution.” Ecology 64: 213–214. 10.2307/1937347.

Souza-Alves, J. P., F. B. Baccaro, I. P. Fontes, M. A. Oliveira, N. M. O. Silva, and A. A. Barnett. 2021. “For Emergency Only: Terrestrial Feeding in Coimbra-Filho’s Titis Reflects Seasonal Arboreal Resource Availability.” Primates 62: 199–206. 10.1007/s10329-020-00859-6.

Souza-Alves, J. P., I. Mourthé, R. R. Hilário, J. C. Bicca-Marques, J. Rehg, C. C. Gestich, A. C. Acero-Murcia, et al. 2019. “Terrestrial Behavior in Titi Monkeys (Callicebus, Cheracebus, and Plecturocebus): Potential Correlates, Patterns, and Differences Between Genera.” International Journal of Primatology 40: 553–572. 10.1007/s10764-019-00105-x.

Stelzner, J. K. 1988. “Thermal Effects on Movement Patterns of Yellow Baboons.” Primates 29: 91–105. 10.1007/BF02380852.

Stelzner, J. K., and G. Hausfater. 1986. “Posture, Microclimate, and Thermoregulation in Yellow Baboons.” Primates 27: 449–463.

Stewart, F. A. 2011. “Brief Communication: Why Sleep in a Nest? Empirical Testing of the Function of Simple Shelters Made by Wild Chimpanzees.” American Journal of Physical Anthropology 146: 313–318. 10.1002/ajpa.21580.

Stewart, F. A., A. K. Piel, J. C. Azkarate, and J. D. Pruetz. 2018. “Savanna Chimpanzees Adjust Sleeping Nest Architecture in Response to Local Weather Conditions.” American Journal of Physical Anthropology 166: 549–562.

Sun, C., B. A. Kaplin, K. A. Kristensen, V. Munyaligoga, J. Mvukiyumwami, K. K. Kajondo, and T. C. Moermond. 1996. “Tree Phenology in a Tropical Montane Forest in Rwanda.” Biotropica 28: 668–681. 10.2307/2389053.

Tabuse, Y. 2026. “Behavioral Thermoregulation in Relation to Humidity in Wild Japanese Macaques (*Macaca fuscata yakui*): The Significance of Semi-Shade.” Primates 67: 403–414. 10.1007/s10329-026-01261-4.

Takemoto, H. 2004. “Seasonal Change in Terrestriality of Chimpanzees in Relation to Microclimate in the Tropical Forest.” American Journal of Physical Anthropology 124: 81–92. 10.1002/ajpa.10342.

Terrien, J., M. Perret, and F. Aujard. 2011. “Behavioral Thermoregulation in Mammals: A Review.” Frontiers in Bioscience 16: 1428–1444.

Thompson, C. L., and E. A. Hermann. 2024. “Behavioral Thermoregulation in Primates: A Review of Literature and Future Avenues.” American Journal of Primatology 86: e23614. 10.1002/ajp.23614.

Thompson, C. L., S. H. Williams, K. E. Glander, and C. J. Vinyard. 2016. “Measuring Microhabitat Temperature in Arboreal Primates: A Comparison of On-Animal and Stationary Approaches.” International Journal of Primatology 37: 495–517. 10.1007/s10764-016-9917-x.

van Lawick-Goodall, J. 1968. “The Behaviour of Free-Living Chimpanzees in the Gombe Stream Reserve.” Animal Behaviour Monographs 1: 161–IN12. 10.1016/S0066-1856(68)80003-2.

Wickham, H., M. Averick, J. Bryan, W. Chang, L. D. McGowan, R. François, G. Grolemund, et al. 2019. “Welcome to the tidyverse.” Journal of Open Source Software 4, no. 43: 1686. 10.21105/joss.01686.

Williamson, E. A., F. G. Maisels, C. P. Groves, B. Fruth, T. H. Humle, F. B. Morton, M. C. Richardson, A. Russon, and I. Singleton. 2013. “Hominidae.” In Handbook of the Mammals of the World, Vol. 3: Primates, edited by R. A. Mittermeier, A. B. Rylands, and D. E. Wilson, 792–854. Barcelona: Lynx Edicions.

Zheng, J., K. Zhang, J. Liang, Y. Li, and Z. Huang. 2021. “Food Availability, Temperature, and Day Length Drive Seasonal Variations in the Positional Behavior of White-Headed Langurs in the Limestone Forests of Southwest Guangxi, China.” Ecology and Evolution 11: 14857–14872. 10.1002/ece3.8171.

Zhou, J., W.-B. Li, X. Wang, and J.-H. Li. 2022. “Seasonal Change in Activity Rhythms and Time Budgets of Tibetan Macaques.” Biology 11: 1260. 10.3390/biology11091260.

